# Altered Gene Regulatory Networks are Associated with the Transition from C_3_ to Crassulacean Acid Metabolism in *Erycina* (Oncidiinae: Orchidaceae)

**DOI:** 10.1101/431460

**Authors:** Karolina Heyduk, Michelle Hwang, Victor A. Albert, Katia Silvera, Tianying Lan, Kimberly M. Farr, Tien-Hao Chang, Ming-Tsair Chan, Klaus Winter, Jim Leebens-Mack

## Abstract

Crassulacean acid metabolism (CAM) photosynthesis is a modification of the core C_3_ photosynthetic pathway that improves the ability of plants to assimilate carbon in water-limited environments. CAM plants fix CO_2_ mostly at night, when transpiration rates are low. All of the CAM pathway genes exist in ancestral C_3_ species, but the timing and magnitude of expression are greatly altered between C_3_ and CAM species. Understanding these regulatory changes is key to elucidating the mechanism by which CAM evolved from C_3_. Here we use two closely related species in the Orchidaceae, *Erycina pusilla* (CAM) and *Erycina crista-galli* (C_3_), to conduct comparative transcriptomic analyses across multiple time points. Clustering of genes with expression variation across the diel cycle revealed some canonical CAM pathway genes similarly expressed in both species, regardless of photosynthetic pathway. However, gene network construction indicated that 149 gene families had significant differences in network connectivity and were further explored for these functional enrichments. Genes involved in light sensing and ABA signaling were some of the most differently connected genes between the C_3_ and CAM *Erycina* species, in agreement with the contrasting diel patterns of stomatal conductance in C_3_ and CAM plants. Our results suggest changes to transcriptional cascades are important for the transition from C_3_ to CAM photosynthesis in *Erycina*.

## Introduction

Crassulacean acid metabolism (CAM) is a carbon concentrating mechanism that evolved multiple times in response to CO_2_ limitation caused by water stress. In C_3_ species, stomata remain open during the day to assimilate atmospheric CO_2_, but water limitation can force stomata to close, resulting in impaired CO_2_ fixation at the expense of growth. When water stress is prolonged, stomatal closure in C_3_ plants can become debilitating. CAM species circumvent prolonged stomatal closure by opening stomata at night and fix CO_2_ nocturnally, when evapotranspiration rates are on average lower. CO_2_ is temporarily stored as malic acid in the vacuoles until day time, when stomata close and malic acid is moved back into the cytosol for decarboxylation. The resulting increase of CO_2_ levels near ribuslose-1,5-bisphosphate carboxylase/oxygenase (RuBisCO) results in highly efficient CO_2_ reduction via C_3_ photosynthesis CAM is associated with a number of anatomical, physiological and genetic change, including alterations to leaf anatomy (Nelson and Sage, 2008; Zambrano et al., 2014), stomatal opening at night, and tight regulation of metabolic genes within day/night cycles. Despite the complexity of these evolutionary novelties, CAM plants are found in a wide range of plant families, including eudicot species in the Euphorbiaceae (Horn et al., 2014) and Caryophyllales (Guralnick et al., 1984; Moore et al., 2017; Winter and Holtum, 2011) and monocot lineages in Agavoideae (Abraham et al., 2016; Heyduk et al., 2016), Orchidaceae (Silvera et al., 2009, 2010), and Bromeliaceae (Crayn et al., 2004).

The CAM pathway is well-described biochemically (Holtum et al., 2005) and contemporary genomics approaches are beginning to shed light on the genetic basis of CAM (Abraham et al., 2016; Cushman et al., 2008; Dever et al., 2015) (Fig. 1). As CO_2_ enters the chloroplast-containing cells as night, it is initially converted to HCO_3_ ^-^ facilitated by a carbonic anhydrase (CA). HCO_3_ ^-^ is then fixed by phosphoenolpyruvate carboxylase (PEPC) using phosphoenolpyruvate (PEP) as the substrate. Carboxylation of PEP results in oxaloacetate (OAA), which is subsequently converted to malic acid by malate dehydrogenase (MDH). Malic acid is then moved into the vacuole for storage. The vacuolar transporter of malic acid is not known for certain, although previous studies have pointed to aluminum-activated malate transporters (ALMT) as a candidate (Kovermann et al., 2007; Yang et al., 2017). During the day, the malic acid is released from the vacuoles either via a passive process or through as-yet undescribed transporter. The malic acid is then decarboxylated to CO_2_ and PEP using two decarboxylation pathways: NAD and/or NADP malic enzymes together with pyruvate, phosphate dikinase (PPDK), or MDH and phosphoenolpyruvate carboxykinase (PEPCK)

**Figure 1.**
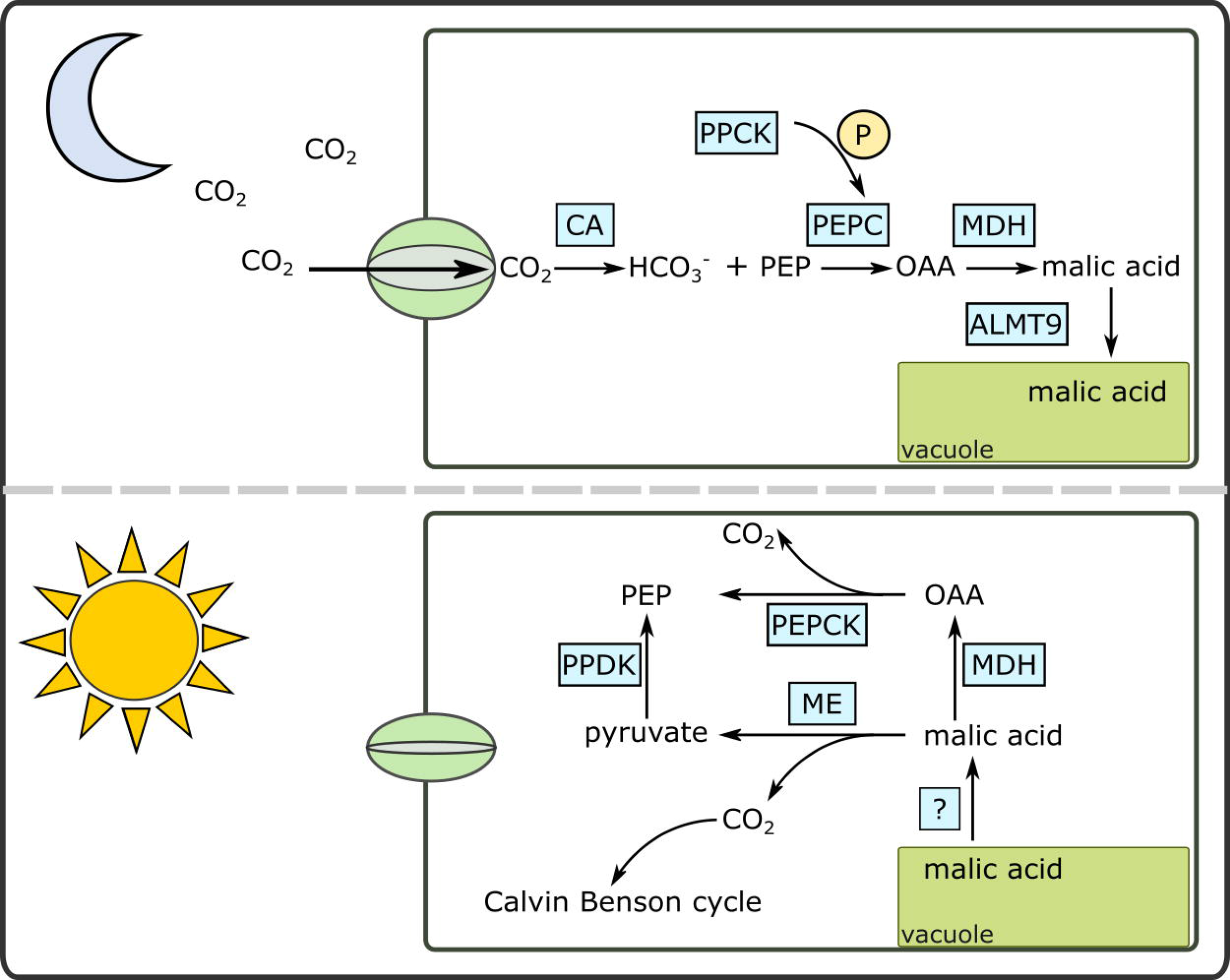
A simplified diagram of the Crassalucean acid metabolism (CAM) pathway under day and night conditions. ALMT9 – aluminum activated malate transporter; CA – carbonic anhydrase; MDH – malate dehydrogenase; OAA – oxaloacetate; ME – malic enzyme (NAD or NADP); PEPC – phosphoenolypyruvate; PEPCK – PEP carboxykinase; PPCK – PEPC kinase; PPDK – pyruvate, phosphate dikinase.

While the roles of canonical CAM enzymes are considered novel in CAM species, they are all present in C_3_ ancestral species as well. As a result, the evolution of CAM likely involved alterations to gene copies, including changes to protein sequences and regulatory motifs. For example, PEPC in C_3_ species can play several roles depending on tissue type and developmental stage, including providing carbon backbones to the citric acid cycle and providing malate for cellular pH balance (Aubry et al., 2011; Winter et al., 2015), but its core function as a carboxylating enzyme remains unchanged in C_3_ and CAM species. Studies of molecular evolution of PEPC in CAM species have largely determined that there are CAM-specific copies of the enzyme that are differentially expressed in C_3_ and CAM species (Gehrig et al., 2001; Lepiniec et al., 1994; Ming et al., 2015; Silvera et al., 2014). In some cases, these CAM-specific PEPC gene copies have been shown to share sequence similarity across closely related but independently derived CAM taxa (Christin et al., 2014; Silvera et al., 2014). Additionally, genomic screens across many canonical CAM gene promoters have revealed an enrichment of circadian clock motifs (Ming et al., 2015), implicating alterations to transcription factor binding sites during CAM evolution. Because of the strong influence of the internal circadian clock on CAM (Hartwell, 2005), we might expect that genes involved in the CAM regulatory pathway should be controlled by a co-expressed circadian master regulator (Borland et al., 1999; Nimmo, 2000; Taybi et al., 2000b).

Elucidation of regulatory changes requires comparative analysis between closely related C_3_ and CAM species, and this can be accomplished through RNA-Seq analyses. One of the largest plant families with multiple origins of CAM is the Orchidaceae; known for floral diversity and inhabiting a broad range of habitats, the evolution of CAM in predominantly epiphytic lineages may have also contributed to orchid diversity (Silvera et al., 2009). Epiphytic species constitute more than 70% of the Orchidaceae (Chase et al., 2015; Gravendeel et al., 2004), and many exhibit different degrees of CAM (Silvera et al., 2005). In a large proportion of these, CAM is weakly expressed relative to C_3_. Weakly expressed CAM may represent an evolutionary end point, or may be an important intermediate step on the evolutionary path between constitutive C_3_ and constitutive CAM. Because many genera within the Orchidaceae include both C_3_ and weak/strong CAM species, the orchids are an attractive family to study the evolution of CAM photosynthesis. The subtribe Oncidiinae is one of the most diverse subtribes within Orchidaceae and it is part of a large epiphytic subfamily (Epidendroideae) in which CAM may have facilitated the expansion into the epiphytic habitat (Silvera et al., 2009). Despite the prevalence of CAM within the subtribe, the genus *Erycina* is particularly interesting because it has both CAM and C_3_ species. *Erycina pusilla* is a fast-growing CAM species with transformation capability and has the potential to be a model species for studying CAM photosynthesis in monocots (Lee et al., 2015). Comparative investigations of *E. pusilla* and its C_3_ relative, *E. crista-galli*, can therefore offer valuable insight into studying the evolution and regulation of CAM photosynthesis in the Orchidaceae. Through comparative, time-course RNA-Seq analysis of *E. pusilla* and *E. crista-galli*, we aim to understand 1) the changes in expression of core CAM genes between C_3_ and CAM *Erycina* species and 2) which regulatory changes are required for the evolution of CAM.

## Materials and Methods

### Plant growth and RNA-Seq tissue collection

*Erycina pusilla* (L.) N.H.Williams & M.W.Chase (CAM) seedlings were cultivated on solid PSYP medium comprising 2 g/L Hyponex No. 1, 2 g/L tryptone, 20 g/L sucrose, 0.1 g/L citric acid, and 1 g/L active charcoal in flasks. The pH of the medium is adjusted to 5.4 before autoclaving and gelling with 3 g/L Phytagel. Plants were grown in 12-hour day and 12-hour night conditions over three independent dates in a growth chamber at the University at Buffalo, with temperatures set to 22-25C and lights on at 6 am for a 12 hour photoperiod. Light intensity was between 95-110 μmol m^−2^ s^−1^. Leaf samples for RNA-sequencing were collected every 4 hours directly from plants grown on sealed flasks for the first two experiments (January and February 2015, Set 1 and Set 2 respectively) and every 2 hours from the final experiment (October 2015, Set 3), where both medium-sized and large plants were collected. Because *Erycina* species are considered miniatures and are therefore relatively small for destructive leaf sampling, we use individual genets as biological replicates at each time point. Leaf samples were flash frozen in liquid N_2_ and stored at −80°C.

*Erycina crista-galli* (Rchb.f.) N.H.Williams & M.W,Chase (C_3_) plants were wild-collected from Peña Blanca, District of Capira, Republic of Panama at 858m above sea level, then grown and propagated in a commercial orchid greenhouse in Bajo Bonito, District of Capira, Republic of Panama. Plants were fertilized once a week week alternatively with a 20-20-20 or 16-32-16 N-P-K fertilizer. Similarly sized and aged plants were moved into an environmental growth chamber at the Smithsonian Tropical Research Institute laboratories (Panama City, Panama) in April 2016, where they were allowed to acclimate for 48 hours to the following conditions: 12 hour light/dark cycle (lights on 6 a.m.), 25°C/22°C day/night temperatures, 60% humidity, and a light intensity of 30 μmol m^−2^ s^−1^, which is similar to the light intensity this species would experience naturally. Biological replicates (consisting of entire shoots without root tissue) were sampled every 4 hours over a 24-hour period, starting at ZT0 (lights on, 6 a.m.) with 4 replicates per time point. Tissue was flash frozen in liquid nitrogen and stored as described above for *E. pusilla*.

RNA was isolated from leaf tissue of both *Erycina* species using the RNeasy Plant Mini Kit (Qiagen). RNA samples were subsequently quantified via Nanodrop and checked for integrity with a Bioanalyzer v2100. RNA libraries were constructed using the Kapa mRNA stranded kit with a combinatorial barcoding scheme (Glenn et al., 2016). Libraries were sequenced on an Illumina NextSeq500 with PE75 reads, pooling 30-32 samples per run. A summary of the data can be found in Supplemental Table 1.

### Gas exchange

Individual shoots (leaves emerging from a common base) of *E. pusilla* and *E. crista-galli* species were individually sealed within a CQP 130 porometer gas-exchange cuvette (Walz, Effeltrich, Germany), located inside an environmental chamber (Environmental Growth Chambers, OH, USA) operating on 12h light (6AM to 6PM) at 28°C, and 12h dark (6PM to 6AM) cycle at 22°C. Light intensity inside the chamber was 230 µmol m^−2^ s^−1^. Plants were watered 3-4 times daily and humidity inside the chamber was maintained at near 60%. Continuous net CO_2_ exchange was measured for each plant for up to 8 day/night cycles with data points obtained every 4 minutes. For *E. pusilla*, water was withheld from one of the three plants measured (drought stress) between the fifth and the eighth day; regarding *E. crista-galli,* water was withheld from the third to the fifth day for one plant of the three measured. Data is presented for all 3 replicates of each of the two *Erycina* species in Supplemental Figure 1.

### Titratable Acidity

*Erycina crista-galli* plants were too small for both transcriptomic and titratable acidity analysis, therefore leaf tissue from this species was collected only for RNA-Seq. Leaf samples from *E. pusilla* were collected from the same plant and at the same time such that half the plant was used for RNA-sequencing and the other half was used for titratable acidity assays. Leaves from mature plants were collected every 4 hours, flash frozen in liquid N_2_, weighed, and boiled in 20% ethanol and deionized water. Titratable acidity was measured as the amount of 0.002M NaOH required to neutralize the extract to a pH of 7. Because leaf tissue was limited for *E. crista-galli* plants, we conducted titrations on three plants that were not sampled for RNAseq to confirm their status as C_3_. Samples were collected in greenhouse conditions at dawn and dusk with three replicates at each time. Titratable acidity was measured as for *E. pusilla* but using 0.001M KOH.

### Transcriptome assembly

An initial *de novo* transcriptome was assembled from Sets 1 and 2 for *E. pusilla* sequences and from all samples sequenced for *E. crista-galli* using Trinity v2.0.6 (Haas et al., 2013). Reads were cleaned using Trimmomatic (Bolger et al., 2014) as implemented in Trinity, and assemblies were made on *in-silico* normalized reads. An initial evaluation of read mapping results from *E. pusilla* Sets 1, 2, and 3 showed a large degree of variation among replicates; to reduce this variation, we used only reads from Sets 1 and 2, as well as medium sized plants from Set 3. All reads for *E. crista-galli* were included in the analysis. These data were further reduced to include only 4 replicates per time points for a total of 24 samples. Replicates were chosen randomly. Read mapping and abundance estimation for transcripts was conducted separately in each species using RSEM v1.3.0 (Li and Dewey, 2011) and Kallisto (Bray et al., 2016). Transcripts with a transcripts per kilobase million mapped (TPM) < 2 were removed and the reads were re-mapped to the filtered assemblies.

### Ortholog circumscription and isoform filtering

To determine gene family circumscription and annotation, all transcripts were sorted into 14 orthogroup gene families from the genomes of the following: *Amborella trichopoda, Ananas comosus, Arabidopsis thaliana, Asparagus officinalis, Brachypodium distachyon, Carica papaya, Dendrobium catenatum*, *Elaeis guineensis*, *Musa acuminata*, *Oryza sativa*, *Phalaenopsis equestris*, *Solanum lycopersicum*, *Sorghum bicolor, Spirodella polyrrhiza*, *Vitis vinifera*, and *Zostera marina*. Assembled transcripts were first used to query the genome database using blastx and sorted to gene families (orthogroups) based on best BLAST hit. The gene families were annotated by *Arabidopsis* members, using TAIR 10 (www.arabidopsis.org) classifications. Transcripts were retained only if they 1) had a length less than the maximum sequence of a gene family based on the sequenced genomes and 2) had a length no less than 50% of the minimum sequence length based on sequenced genome members of that gene family.

Trinity produces both gene components and subsidiary isoforms, which may represent true alternative splice isoforms or allelic or paralogous sequence variants. To mitigate dilution of read mappings to multiple isoforms, we instead used gene components (hereafter referred to as transcripts) for all further analyses (including gene level read mapping from RSEM). For gene tree estimation we took the longest isoform per component per orthogroup, using our minimum/maximum orthogroup filtered data set. Scripts for orthogroup sorting and filtering can be found at www.github.com/kheyduk/Erycina.

### Time-dependent clustering

To incorporate time into our clustering analysis, we used R software package maSigPro v1.46.0 (Conesa et al., 2006; Nueda et al., 2014), which analyzes expression data for patterns across time by fitting each gene’s expression pattern to a polynomial using stepwise regression. Cross-normalized read counts and a negative binomial distribution for the generalized linear models were used. For each transcript, maSigPro estimated up to a 4^th^ degree polynomial and tested the fit via ANOVA. Transcripts that had significantly time-structured expression (Benjamini & Hochberg adjusted p < 0.05) were retained while all others were removed from further analysis. Additionally, any genes considered overly influential based on DFBETAS diagnostic (how much an observation affects the estimate of a regression coefficient) (Belsley et al., 1980) were also removed. In total, 1,515 transcripts from *E. pusilla* and 505 transcripts from *E. crista-galli* were removed as influential genes.

The remaining transcripts that did show time-dependent expression (n=7,066 in *E. pusilla* and n=7,127 in *E. crista-galli*) were clustered by fuzzy clustering based on similarity in expression profiles. An optimal fuzzifier *m*, a parameter that determines how much clusters can overlap, was calculated in the Mfuzz package (Kumar and E Futschik, 2007) of R for each species (*m*=1.09 for both *E. pusilla* and *E. crista-galli*). The number of groups *k* for each species was determined by choosing a value which minimizes the within-group variance (Supplemental Figure 2); a *k* of 6 was used for both *E. pusilla* and *E. crista-galli*. Z-scores of normalized counts were calculated for each gene in each cluster, as well as a median cluster expression, for each species separately.

**Figure 2.**
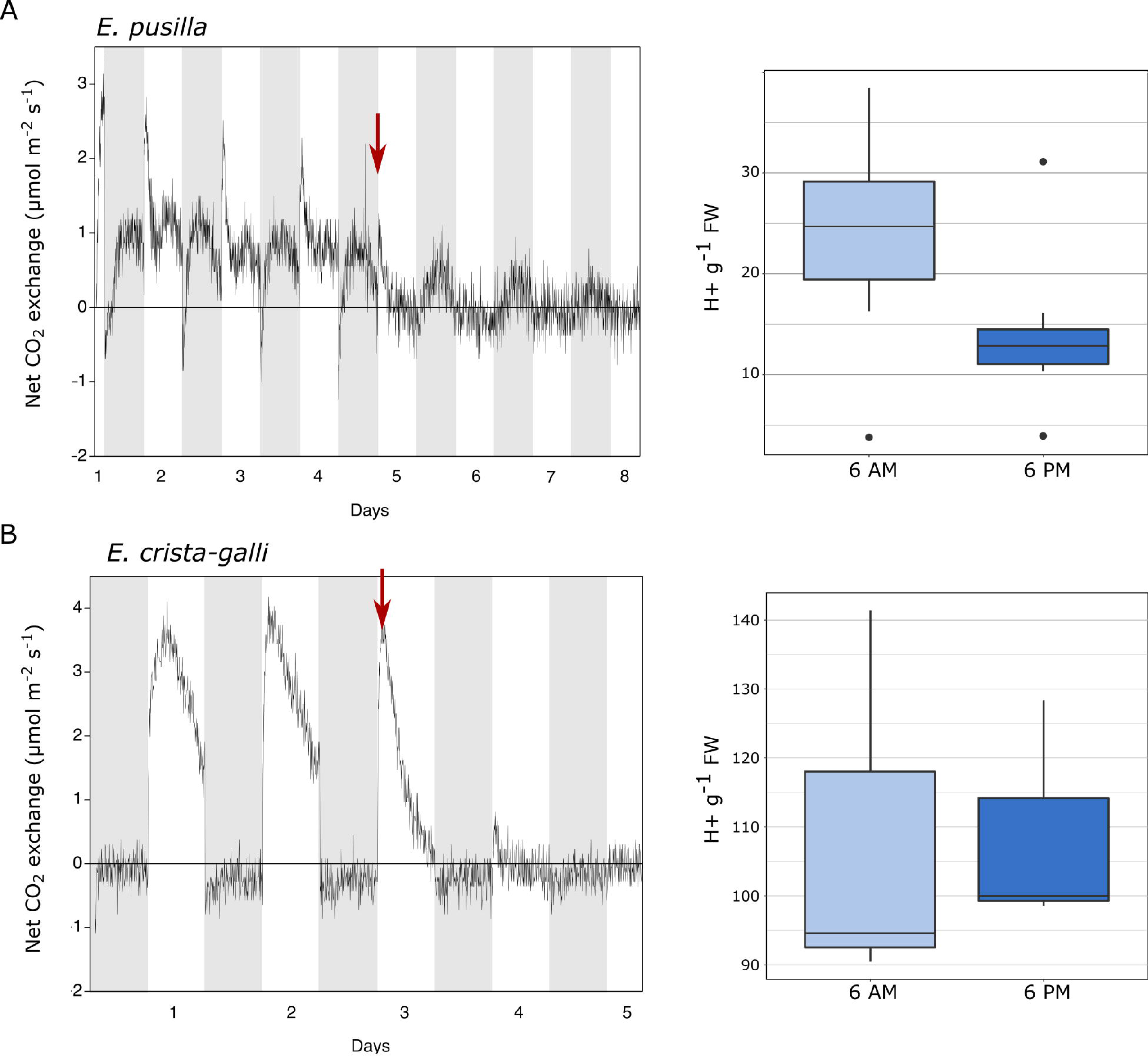
Gas exchange and titratable acidity for A) *Erycina pusilla* and B) *Erycina crista-galli.* Gas exchange is shown for a single plant; replicate plant gas exchange plots can be found in Supplemental Figure 1. Drought induction is indicated with a red arrow. Titrations are shown for dawn and dusk; full titratable acidity values are in Supplemental Table 2.

### Gene trees and expression

Gene trees were estimated for phosphoenolpyruvate carboxylase (PEPC) and its kinase (PPCK) by first aligning nucleotide sequences from *Erycina* transcripts and their associated gene family members from the sequenced genomes using PASTA (Mirarab et al., 2014), then estimating trees using RAxML (Stamatakis, 2006). Gene expression for genes of interest was plotted based on averaged transcripts per million mapped (TPM) for each replicate.

### Network analysis

To identify regulatory candidates possibly involved in CAM and examine the relationships between genes within clusters, we used the ARACNe-AP v1.4 (Lachmann et al., 2016) algorithm to create networks of co-expressed transcripts from both species separately. Briefly, the algorithm randomly samples gene pairs and uses an adaptive partitioning approach to infer a pairwise mutual information (MI) statistic, or measure of statistical dependence, between them. This process is repeated iteratively for a specified number of bootstraps, while at each step removing indirect interactions. A final network is built based on the consensus of all bootstrap runs. Although ARACNe provides an option to specify transcription factors to generate a directed network by only considering interactions with a transcription factor source, we chose to generate an undirected gene co-expression network of genes that were significantly time-structured based on our maSigPro analysis, using 100 bootstrap replicates in ARACNe.

We imported network data into Cytoscape (Shannon et al., 2003) to generate visualizations and calculate network statistics. Nodes were color coded by their cluster membership and scaled to represent number of connections. Network statistics were exported and further analyzed at the orthogroup level. We calculated which orthogroups had the largest average difference in connectivity between the two species. We first calculated the average connectivity (number of directed edges, output from Cytoscape Network Analysis) for each orthogroup per species, then normalized these via Z-scores and subtracted the Z-score of the orthogroup in *E. pusilla* from that in *E. crista-galli*. Outliers – those orthogroups with the largest difference between species in connectivity – had Z-score differences above and below the upper and lower quantiles (Supplemental Figure 3).

**Figure 3.**
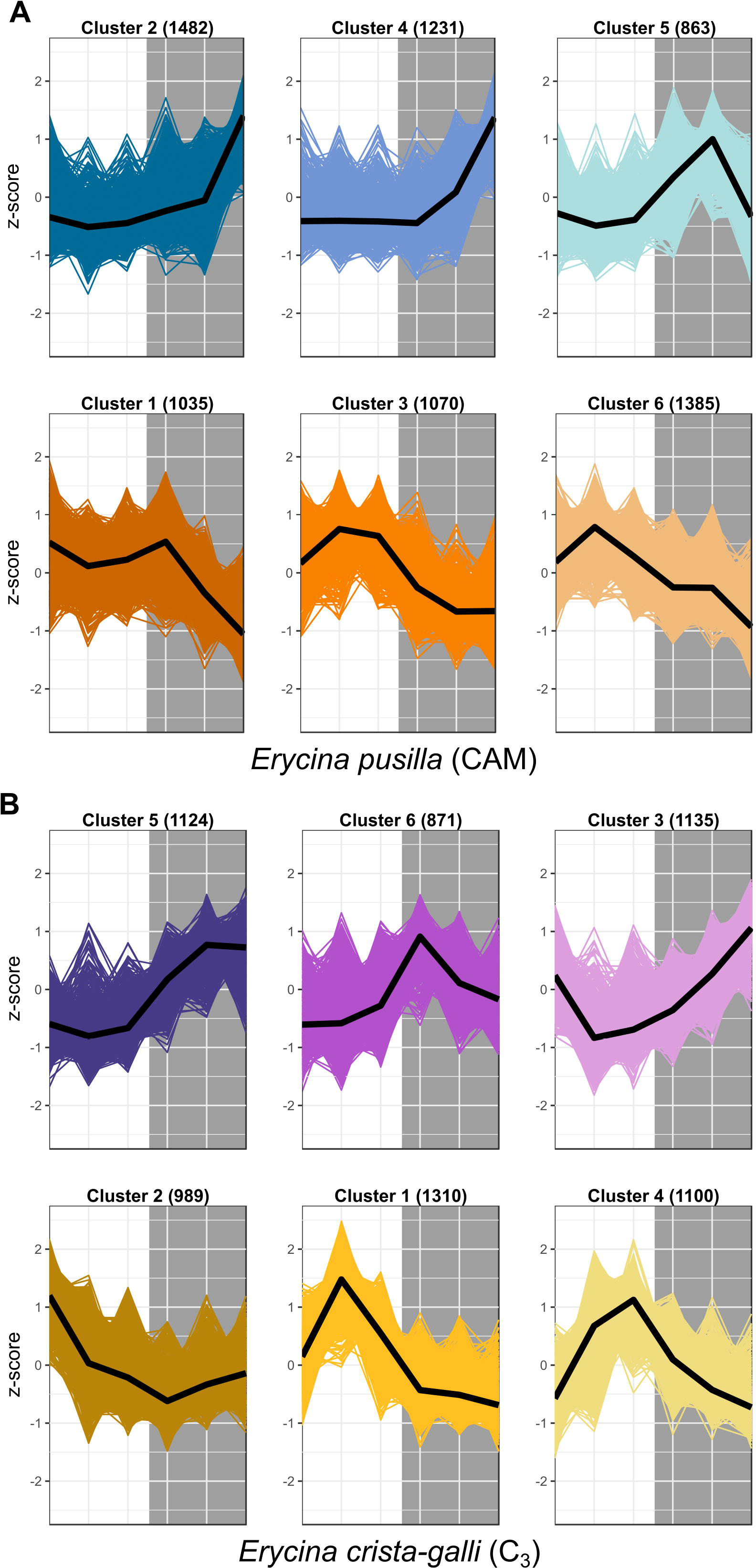
Expression z-scores for each gene in each cluster for both A) *E. pusilla* and B) *E. crista-galli*, with median expression shown in the black line. Clusters represent co-expressed genes within each species’ time-structured transcripts. Cool colors (blue and purple) are clusters with nighttime biased expression, whereas warmer colors (orange and yellow) are clusters whose transcripts increase in expression during the day.

The outlier orthogroups with large changes to connectivity between species (Supplemental Table 3) were explored for genes of interest. The largest difference in connectivity was found in a E3 ubiquitin ligase gene family shown to play a role in ABA signaling, with the *Arabidopsis* homolog known as ring finger of seed longevity1 (RSL1). To explore differences in network connections of RSL1 between the two species, we employed the diffusion algorithm (Carlin et al., 2017) in Cytoscape which finds strongly interactive nodes to a target of interest. For both species, we found the diffusion network for the RSL1 gene. Only a single gene copy of RSL1 was time-structured in *E. crista-galli*, but *E. pusilla* had two copies found in the ARACNe network. One gene copy had only a single connection to any other gene in the network and was not analyzed further. The other copy in *E. pusilla*, which had 72 directed connections, was used as the center of the diffusion network. Diffusion networks were compared for orthogroup content between species using a hypergeometric test. Gene Ontology (GO) terms were compared for the two RSL1 subnetworks and checked for enrichment using hypergeometric test (using all GO terms found in either ARACNe network as the universe), correcting for multiple testing with Bonferroni-Holm significance correction.

## Results

### Gas exchange patterns and titratable acidity

Gas exchange data collected continuously showed net nighttime CO_2_ uptake in CAM *E. pusilla* under both well-watered conditions and while drought stressed (Fig. 2a). C_3_ *Erycina crista-galli* displayed net CO_2_ uptake during the light period only. There was no net uptake of CO_2_ at night. Nonetheless, under drought stress, a slight decrease in respiratory loss of CO_2_ at night may indicate low levels of CAM cycling. Titration data collected from the same plants and at the time of RNA-sampling in *E. pusilla* confirms CAM function in the plants used for gene expression analysis, with a significant increase in leaf titratable acids occurring towards the end of the dark period (Fig. 2A, 6AM), and a reduction in total acids during the day period. Although the C_3_ *E. crista-galli* had higher overall levels of leaf acids, there was no significant diurnal fluctuation (Fig. 2b).

### Clustering of genes with time-structured expression profiles

After filtering by minimum/maximum length each transcript’s orthogroup, 23,596 and 26,437 genes were retained in *E. pusilla* and *E. crista-galli*, respectively. Both species had a similar number of genes that were significantly time-structured (~7,000) according to maSigPro. Each species had best fit to k=6 clusters, with three clusters showing nighttime biased expression and three with daytime bias (Fig. 3). Expression of PEPC, the initial carboxylating enzyme in the CO_2_ fixation pathway at night, increased in expression just before the onset of darkness in the CAM species *E. pusilla* (Fig. 4A). In contrast, there was a low, but significant, time-structured expression pattern of PEPC in the C_3_ species *E. crista-galli* (Fig. 4B). The dedicated kinase, PPCK, which phosphorylates PEPC and allows it to function in the presence of malate, likewise showed a strong nocturnal increase in expression in the CAM species, with similar levels of expression in the C_3_ species (Fig. 4B).

**Figure 4.**
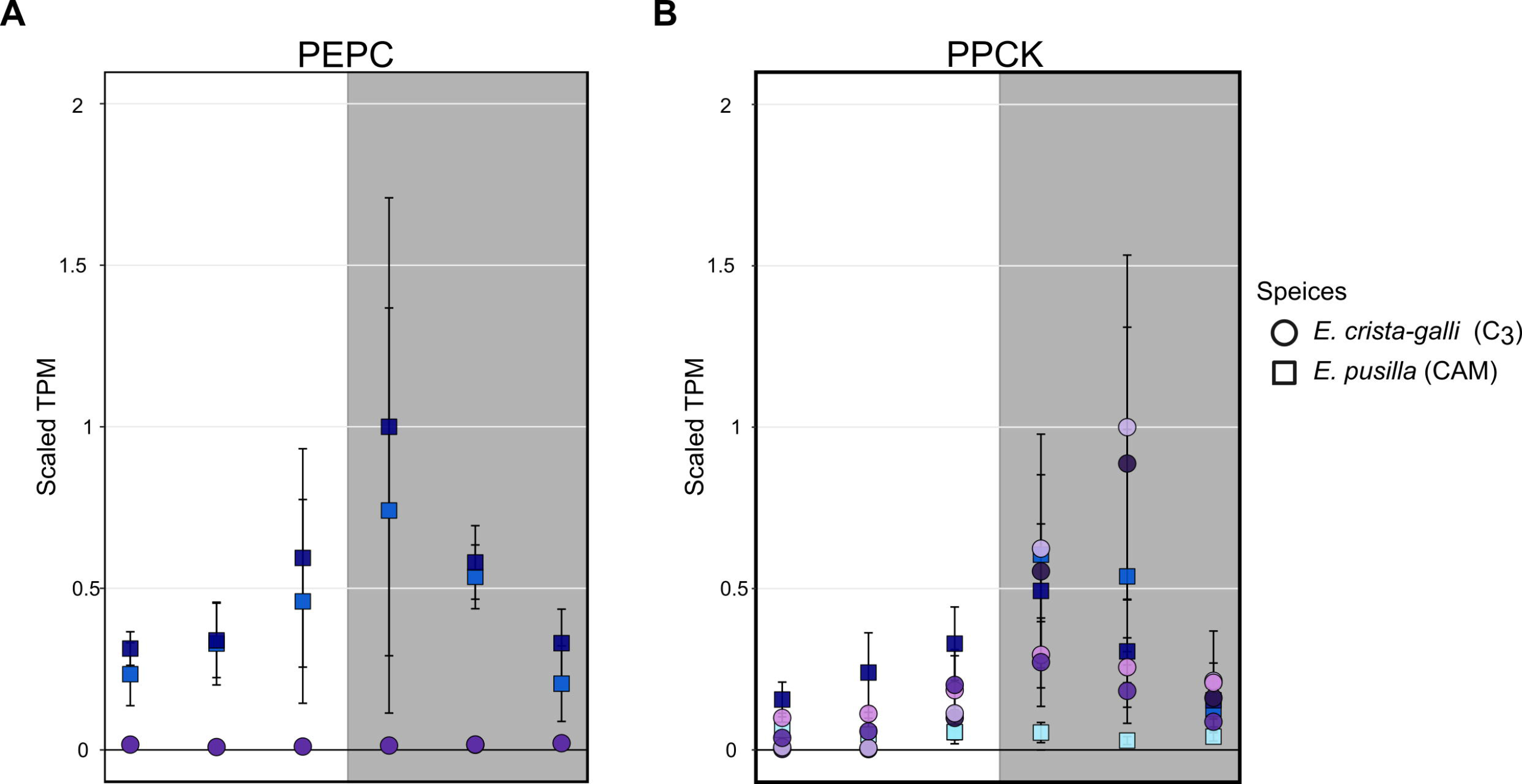
Expression of A) phosphoenolpyruvate carboxylase (PEPC) and B) PEPC kinase in *E. pusilla* (squares, blue tones) and *E. crista-galli* (circles, purple tones). Different shades of color represent different assembled gene copies, and only transcripts that were found to be significantly time-structured are shown. Average TPM and standard deviation, scaled to the maximum mean TPM value across all copies and species, is plotted for all 6 time points, with the grey background indicating nighttime samples.

### Network Comparisons

The network for C_3_ *E. crista-galli* had 119,338 directed connections between 4,828 nodes, whereas the CAM *E. pusilla* network was notably less connected, with only 76,071 connections between 4,591 nodes. Although the number of genes in each network was similar, overall connectivity of *E. pusilla* is easily seen in both the fewer number of connections as well as the mean number of connections (34.3 in *E. pusilla* vs. 50.3 in *E. crista-galli*). As expected, genes from the same co-expressed cluster (Fig. 3) were grouped within the larger ARACNe network for each species (Fig. 5A,C).

**Figure 5.**
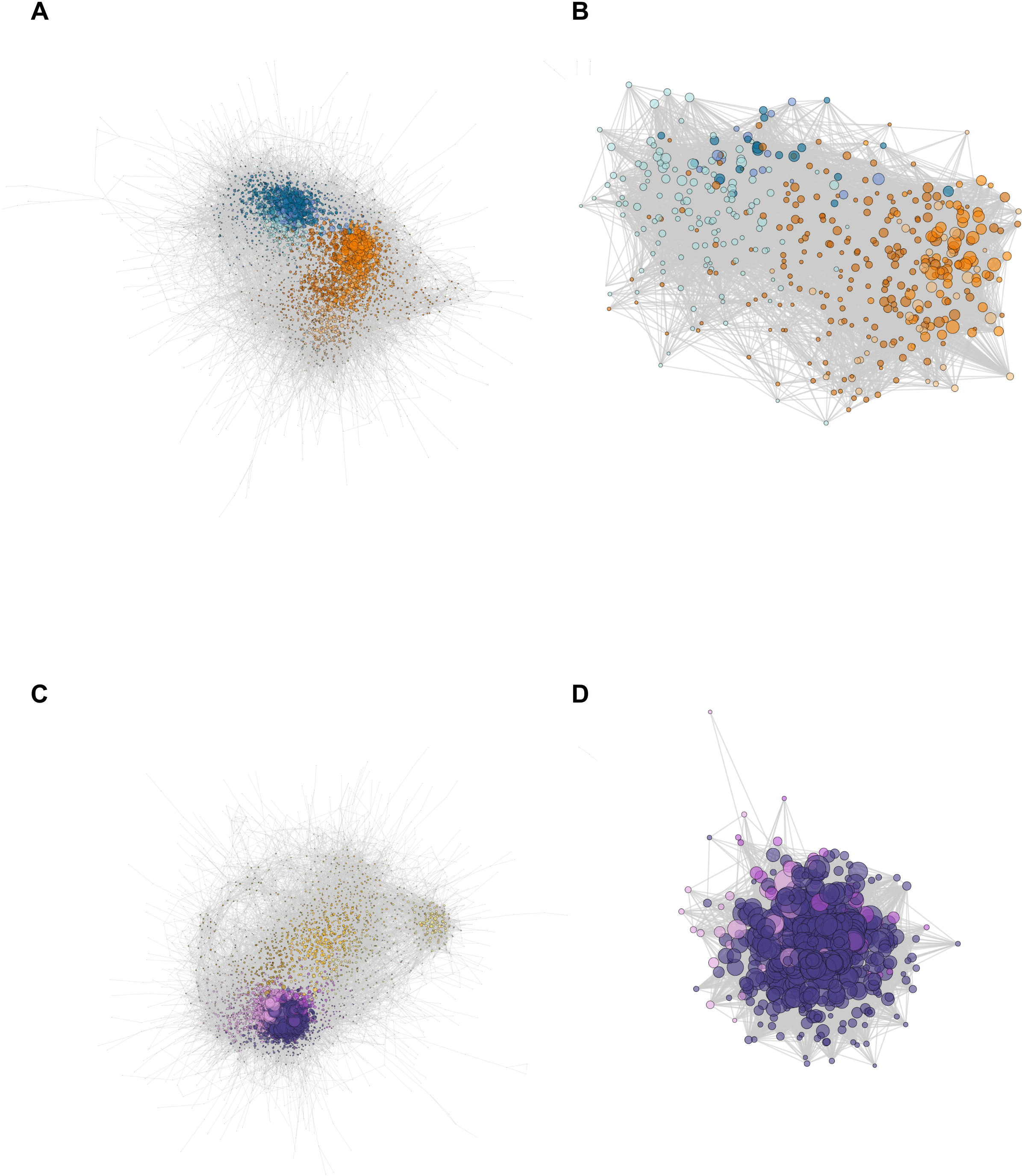
Network constructions of A) *E. pusilla* and C) *E. crista-galli* based on ARACNe network analysis, and the ring finger seed longevity1 (RSL1) subnetworks for B) *E. pusilla* and D) *E. crista-galli* calculated via the diffusion algorithm in Cytoscape. Colors correspond to Figure 3 cluster colors – cooler colors (blues in A and B and purples in C and D) are clusters that have an increase in expression at night and warmer colors (oranges in A and B and yellows in C and D) are clusters that have increases in expression during the day. Dots represent genes, scaled by the number of edges (maximum number of directed edges=435).

Comparison of network connectivity of the time-structured genes found 149 outlier transcripts that had large species differences in the number of connected directed edges (Additional file 4); these were largely skewed toward increased connectivity in *E. crista-galli* (n=90). Annotations of these outliers revealed a number of genes involved in stomatal opening/closing and ABA signaling. GO term enrichment indicates that outliers that skew toward more connectivity in C_3_ *E. crista-galli* were enriched for vacuolar and tonoplast membrane proteins and potassium and calcium transport. Genes that were more connected in CAM *E. pusilla* were enriched for genes involved in aldehyde dehydrogenase activity, among other functions (Supplemental Table 4).

A E3 ubiquitin ligase also known as ring finger of seed longevity1 (RSL1) had the greatest difference in connectivity between the two species. RSL1 has been shown to be a negative regulator of ABA signaling (Bueso et al., 2014) and was chosen as a center node for comparison between the two species. The diffusion algorithm used to create subnetworks defaults to producing a subnetwork that is 10% of the total nodes in the larger network; as a result, both species subnetworks were roughly the same size, containing about 400 genes. However, the connectivity of those subnetworks differed greatly (Fig. 5B,D); the C_3_ *E. crista-galli* RSL1 subnetwork contained 30,392 connections, whereas the CAM *E. pusilla* network had only 9,244. The subnetworks differed in their gene content as well. There contained 429 and 427 orthogroups in *E. pusilla* and *E. crista-galli,* respectively, but only 57 orthogroups were shared between the two (this was not significantly under-enriched via a hypergeometric test (p=1)). While all of the genes in the *E. crista-galli* RSL1 subnetwork were in night-biased expression clusters (485), *E. pusilla* had more subnetwork genes in day-biased clusters (298) than in night biased ones (162). GO term enrichment indicates both subnetworks are enriched for chloroplast, chloroplast stroma, and photosynthesis (Supplemental Table 5).

While both subnetworks were centered on RSL1 and generally were enriched for similar types of genes involved in chloroplast functions and photosynthesis, there were substantial differences in gene content between the networks. The focal gene RSL1 acts as a master negative regulator of ABA signaling pathway by targeting pyrabactin resistance 1 (PYR1) and PYR-like (PYL) ABA receptors for degradation. More generally its role in protein ubiquitination is relatively unknown. RSL1 was the third most connected node in the C_3_ *E. crista-galli* subnetwork (419 directed connections; most connected node had 423 connections). In CAM *E. pusilla*, RSL1 was 173^rd^ out of 460 genes in connectivity. *E. pusilla* had a number of ABA responsive genes in its RSL1 subnetwork that *E. crista-galli* did not, including protein phosphatase 2C (PP2C), a gene encoding a member of the Snf1-related kinase family, plus a homolog of *ABA Overly Sensitive 5*. Recent work has shown that ABA responses in stomata including *PYR/PYL* and downstream genes are responsible for constitutive stomatal aperture, as well as stomatal responses to drought stress (Gonzalez-Guzman et al., 2012).

*Erycina crista-galli*, on the other hand, had a number of light sensing and circadian clock genes that CAM *E. pusilla* did not have in the RSL1 subnetwork, including *lov kelch protein 2* (LKP2/ZTL gene family), *time for coffee* (*TIC*), *phytochrome B/D* (*PHYB*), and *protein phosphatase 2A* (*PP2A*). LKP2/ZTL and PHYB are both involved in light sensing (blue and red/far-red, respectively). Specifically, *LKP2/ZTL* genes are thought to regulate light induced protein degradation via their function as E3 ligases (Demarsy and Fankhauser, 2009; Ito et al., 2012), and *ztl* mutants in *Arabidopsis* show a prolonged clock period under constant light due to the lack of degradation of clock components via ZTL (Más et al., 2003; Somers et al., 2000). TIC has been found to be responsible for the amplitude of the circadian clock but is not thought to be directly involved in light signaling (Hall et al., 2003). TIC is also implicated in daytime transcriptional induction via its association with the central circadian oscillator late elongated hypocotyl (LHY) (Ding et al., 2007). Finally, PP2A is a member of a large family of plant phosphoprotein phosphatases (PPP) with several cellular roles (Uhrig et al., 2013). PP2A specifically has been implicated in brassinosteroid signaling, light signaling via dephosphorylation of phototropin2 (Tseng and Briggs, 2010), flowering time control (Kim et al., 2002), as well as the induction of CAM under certain abiotic stresses (Cushman and Bohnert, 1999).

Although the C_3_ *E. crista-galli* RSL1 subnetwork contained circadian and light sensing transcripts that were absent in the *E. pusilla* RSL1 subnetwork, these transcripts largely did not have different expression patterns between the two species with the exception of *PHYB* (Fig. 6B). The contrasting gene content of the RSL1 subnetwork suggests light sensing and circadian regulatory cascades comprise large differences between C_3_ and CAM species, rather than levels of gene expression, which were quite similar.

**Figure 6.**
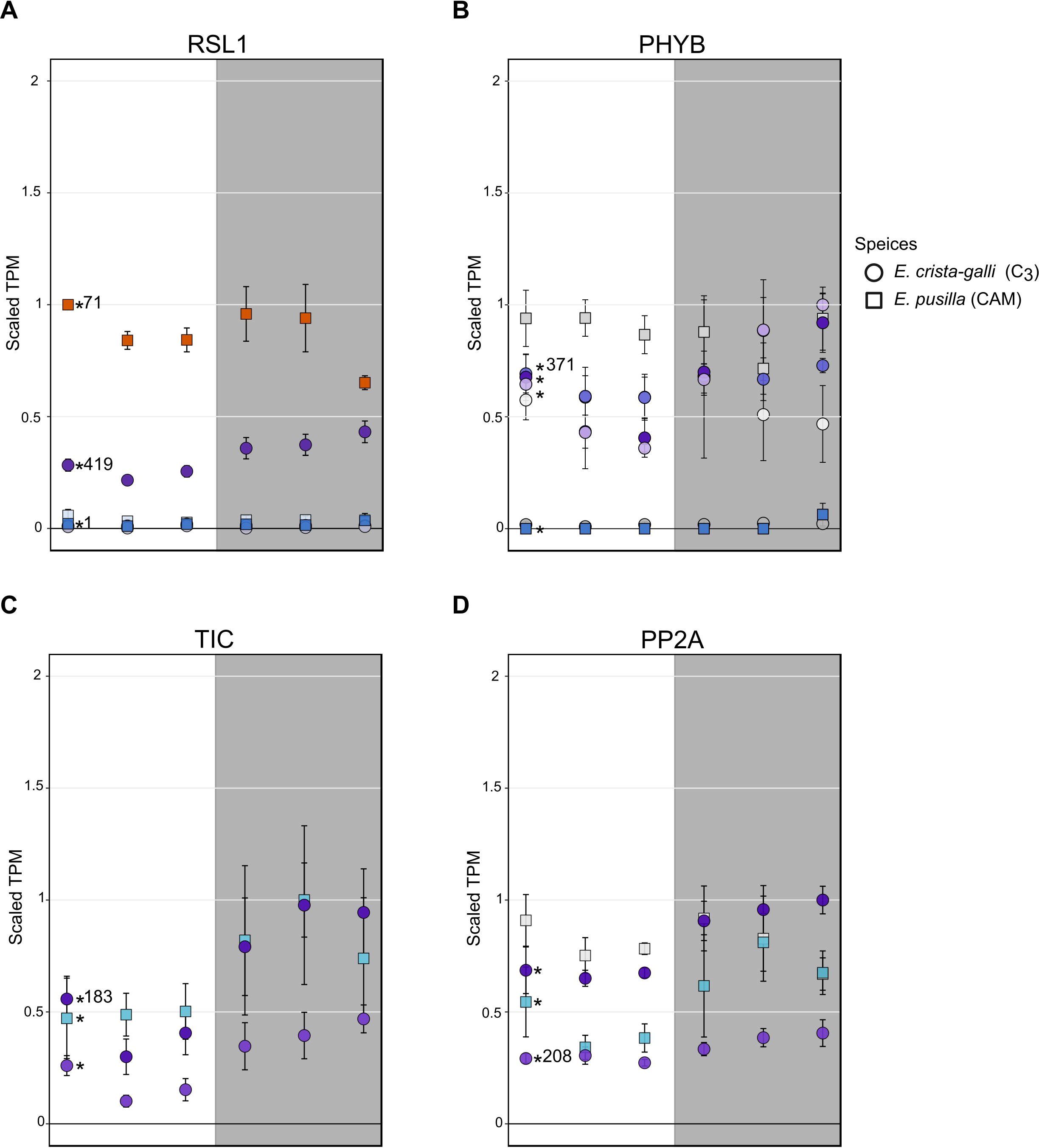
Expression of A) ring finger of seed longevity 1 (RSL1), B) phytochrome B/D (PHYB) C) time for coffee (TIC), and D) protein phosphatase 2A in *E. pusilla* (squares) and *E. crista-galli* (circles). Points are colored by which category of cluster they belong to: blues and purples are genes with increases in expression in the dark in *E. pusilla* and *E. crista-galli*, respectively, while oranges and yellows are genes with increases in daytime expression in *E. pusilla* and *E. crista-galli*, respectively (see Fig. 3). Time-structured genes are marked with an asterisk, while those that were not significantly time-structured are shown in greys. Number of directed edges per gene is shown if they belonged to the RSL1-subnetwork. Average TPM and standard deviation, scaled to the maximum mean TPM value across all copies and species, is plotted for all 6 time points, with the grey background indicating nighttime samples.

## Discussion

### Shared gene expression patterns

While traditionally it was thought that canonical CAM genes should have large differences in timing and magnitude of expression between C_3_ and CAM species, recent work has highlighted that between closely related C_3_ and CAM species, that may not always be the case (Heyduk et al., 2018b). In *Erycina*, it appears that a similar pattern holds. Both the CAM and C_3_ species have time-structured expression of PEPC albeit at very different expression levels. While this alone says little about PEPC’s function in both species, the high nocturnal expression of PEPC’s dedicated kinase, PPCK, in both *E. pusilla* and *E. crista-galli* suggests that PEPC is being phosphorylated in both species and therefore has the need to function in the presence of malate.

It is worth noting that all C_3_ species have the genes involved in the CAM cycle. Many, including PEPC, function in anaplerotic reactions of the TCA cycle. PEPC has also been shown to have a role in malate production for osmotic regulation of stomatal aperture and in CO_2_ fixation in guard cells of tobacco (Asai et al., 2000; Daloso et al., 2015). Because most RNA-seq or gene expression studies to date in C_3_ species sample during the day, understanding how common nocturnal *PEPC* expression is across flowering plants will require more nighttime gene expression studies in C_3_ species, especially in lineages closely related to CAM species. While nearly all the genes in the CAM CO_2_ fixation pathway have known functions in C_3_ species, *PPCK* is a notable exception. Phosphorylation of PEPC by PPCK in CAM and C_4_ species is well-described (Jiao and Chollet, 1989; Nimmo et al., 1986; Taybi et al., 2000a); malate and other organic acids act as negative regulators of PEPC, but phosphorylation of PEPC renders it immune to these negative effects. Thus, phosphorylated PEPC via PPCK is required for high levels of CAM and C_4_ malic acid production. In C_3_ species, however, there is no clear need for heavily phosphorylated PEPC, especially at night, making the expression of *PEPC* and *PPCK* in C_3_ *E. crista-galli* intriguing (though see Sullivan et al., 2004 and Fukayama et al., 2006 for work on nocturnal *PPCK* expression in soybean and rice, respectively).

### Alterations in regulatory pathways between C3 and CAM orchids

Network analysis of co-expressed genes and subsequent comparisons between species can give insights not only into the changes in expression, but also the degree to which a given gene changes in connectivity between species. Extensive connectivity for a gene has long thought to be a signal of a “hub” or master regulatory gene – one that cannot experience large changes in timing or magnitude of expression without significant perturbations to the entire network. A major assumption of co-expression networks is that they rely on mRNA as an accurate predictor of downstream processes; while this is not always the case, recent work showed that although mRNA and protein networks differed in their gene content, they overlapped in gene ontologies and were predictive of pathway regulation in maize (Walley et al., 2016). Additionally, it has been shown that a correlation exists between connectivity of a gene within a network and the evolutionary conservation of the gene’s sequence across a number of flowering plant species (Masalia et al., 2017). It is therefore somewhat surprising to observe that as many as 149 gene families have large changes in connectivity between the two closely related *Erycina* species.

The 149 outlier gene families in *Erycina* were enriched for functions in protein degradation via phosphorylation and ubiquitination. While typically differences in phenotype are considered the result of changes in the abundance of gene products, our data highlight the importance of considering protein degradation as well. The differences in connectivity of protein degradation pathway genes were unbiased between species – in other words, genes involved in protein degradation have increased connectivity in both species. More interestingly, genes that had increased connectivity in C_3_ *E. crista-galli* were enriched for GO terms involved in potassium and chloride channels and membrane proteins associated with chloroplasts and vacuoles. Greater connectivity of such genes in the C_3_ species indicates an increased reliance on ion and metabolite fluxes. In stomatal guard cells, which make up a smaller portion of the whole-leaf transcriptome, these fluxes directly affect stomatal aperture and may play a role in alternative regulation of stomatal opening in C_3_ and CAM species.

### Regulatory changes in ABA, light, and clock perception

Stomatal opening in CAM species has been vastly understudied, despite the opportunities it presents for understanding a fundamental biological process. In C_3_ species, stomatal opening is thought to be regulated by blue and red light inputs, whereas stomatal closing is driven by efflux of potassium cations. It remains largely unknown how stomata sense darkness, but experimental data have suggested a large role of CO_2_ concentrations on stomatal aperture (Cockburn et al., 1979). Draw down of CO_2_ concentrations at night in the intercellular airspace would result in stomatal opening, whereas high concentrations of CO_2_ from decarboxylation during the day may promote stomatal closure. CO_2_ concentrations undoubtedly play some role in the inverted stomatal aperture of CAM species, but more contemporary genomic work has implicated additional levels of regulation in CAM stomata (Abraham et al., 2016; Cushman and Bohnert, 1997; Wai et al., 2017). For example, ABA may be one signaling molecule for nocturnal stomatal closure (Desikan et al., 2004; Gonzalez-Guzman et al., 2012; Merilo et al., 2013). Gene expression results coupled with network analysis in *Erycina* indicate that both ABA signaling and light sensing are likely to be altered. Indeed, the gene with the largest difference in connectivity between species is *RSL1*, which encodes a E3 ubiquitin ligase known to function in stomatal response to ABA. Expression of RSL1 is higher in CAM *E. pusilla* and is clustered with day-biased genes (Fig. 6A), whereas in C_3_ *E. crista-galli* expression is about half that in *E. pusilla* and slightly increases in expression during the dark period. These expression patterns are consistent with stomatal regulation between C_3_ and CAM species. In *E. crista-galli* nighttime ABA may play a larger role in drought-induced or nighttime stomatal closure than in the CAM *E. pusilla,* where the nocturnal stomatal closure driven by ABA signaling must be repressed to allow for nighttime CO_2_ acquisition. It is also unknown what causes daytime stomatal closure in CAM species; while high intracellular CO_2_ concentrations may play a role, so might ABA signaling. *RSL1*’s high connectivity to other genes in the subnetwork of *E. crista-galli* suggests that alterations to regulatory networks are also important for fully functional CAM.

Stomata, while highly responsive to ABA, are also strongly regulated by light inputs. The gene encoding phytochrome B (phyB), a photoreceptor that both regulates transcriptional responses to red and far-red light as well as entrains the circadian clock (Goosey et al., 1997; Más et al., 2000; Ni et al., 1999), was differentially regulated and expressed in the two *Erycina* species (Fig. 6B). While both species had copies of *phyB* that showed time-structured expression, only *E. crista-galli* had a copy that was in the same network as *RSL1* and had many connections to other genes in the same network. A single copy of *phyB* was time-structured in CAM *E. pusilla*, and the expression level relative to copies in C_3_ *E. crista-galli* was quite low. Instead, the constitutively expressed copy of *phyB* in *E. pusilla* had the highest expression, but what it’s role might be given constant expression across time is unclear. The stark difference in both connectivity and expression levels of *phyB* in *E. pusilla* and *E. crista-galli* suggests that light-induced transcriptional regulation has a greater role in the C_3_ species, and that phyB mediated transcriptional regulation in CAM species may be significantly reduced. Additionally, the presence of *PP2A* in the C_3_ *E. crista-galli* RSL1 subnetwork, but not in *E. pusilla,* further indicates that light induced responses are differentially regulated (although expression was similar between the two species, Fig. 6D). PP2A is, among many other tasks, responsible for dephosphorylation of photoropin 2, which subsequently promotes stomatal opening (Tseng and Briggs, 2010; Uhrig et al., 2013).

Differences in light input sensing and signaling between C_3_ and CAM species is not surprising, but relatively little work has focused on this aspect of CAM biology. Previous work assessed light responses in the facultative CAM species *Portulacaria afra* (Lee and Assmann, 1992) and *Mesembryanthemum crystallinum* (Tallman et al., 1997) and showed that both species had reduced stomatal response to blue light signals when relying on the CAM cycle for carbon fixation (though see Ceusters et al., 2014). Additionally, *M. crystallinum* had reduced guard cell zeaxanthin production during the day in the CAM state compared to the C_3_ state. The reduction in zeaxanthin in the CAM state was shown to be a result of the downregulation of the pathway, rather than an aberration in guard cell chloroplasts. Changes to regulation and expression of *phyB* in *Erycina* further support significantly altered light-induced pathways in CAM species relative to C_3_.

*TIME FOR COFFEE* (*TIC*) also stood out in its altered connectivity between the C_3_ and CAM species. In C_3_ *E. crista-galli*, TIC was present in the RSL1 diffusion network whereas it was not for *E. pusilla*. TIC has been shown to be involved in maintaining the amplitude of the circadian clock in *Arabidopsis* (Ding et al., 2007; Hall et al., 2003), as well as regulating metabolic homeostasis and response to environmental cues (Sanchez-Villarreal et al., 2013). *tic* mutants in *Arabidopsis* showed large phenotypic effects ranging from late flowering to anatomical abnormalities. The *tic* mutants also showed extreme tolerance to drought, likely due to increased amounts of osmolytes such as proline and myo-inositol, as well as an accumulation of starch. As a result of the pleiotropic effects of TIC, altered network status of TIC in CAM *E. pusilla* compared to C3 *E. crista-galli* likely results a complex alteration in phenotype. In general, the mechanism that link the circadian clock and CAM photosynthesis are unknown (Boxall et al., 2005), and research to uncover circadian regulation of CAM is limited to transcriptomic studies. Gene expression comparisons between CAM *Agave* and C_3_ *Arabidopsis* revealed changes to the timing of expression of *REVEILLE 1,* a clock output gene that integrates the circadian network to metabolic activity (Yin et al., 2018). While it’s possible that changes to the timing of *REVEILLE 1* are required for CAM evolution, expression differences may also be a result of lineage-specific changes to expression unrelated to CAM. Comparisons of closely related C3 and CAM species of *Erycina* suggest that alteration of transcriptional cascades from circadian oscillators may play a role in the evolution of CAM, rather than large scale changes to the timing or abundance of expression, but additional work to link clock outputs to the CAM phenotype are necessary.

### Implications for the evolution of CAM in Oncidiinae

Both CAM and C_4_ photosynthesis have evolved multiple times across the flowering plant phylogeny, suggesting that the evolution of these complex traits cannot be insurmountably difficult. Recent physiological work demonstrates that the evolution of anatomical traits required for CAM or C_4_ often predates the emergence of strong, constitutive carbon concentrating mechanisms (Christin et al., 2013; Heyduk et al., 2016). Other transcriptomic work has shown that closely related C_3_ and CAM species share the expression of canonical CAM genes, especially *PEPC* and *PPCK* as seen here in *Erycina* (Heyduk et al., 2018b). It has even been suggested that many C_3_ plants already have the nocturnal CAM cycle in place for fixation of respired CO_2_ and generation of amino acids (Bräutigam et al., 2017), but this alone is unlikely to entirely explain the repeated and relatively frequent emergence of CAM on the angiosperm phylogeny.

Indeed it appears that gene expression alone would not facilitate the large-scale transition from C_3_ to CAM. Recent comparative work across multiple C_3_ and C_4_ transcriptomes highlighted the recurrent co-option of highly expressed genes from C_3_ species into C_4_ (Moreno-Villena et al., 2018). The initial co-option of highly expressed gene copies happened early in the evolutionary trajectory between C_3_ and C_4_, and later steps included the refinement of C_3_ enzymes, including kinetic and tissue specificity. It is quite likely that a similar model of evolution holds for CAM, in that C_3_ relatives of CAM lineages have been shown to have similar expression patterns of canonical CAM genes (though, it is worth noting, they are not typically highly expressed in the C_3_ species). In *Erycina*, both C_3_ and CAM species share similar expression profiles for *PEPC* and *PPCK*, with the latter having nearly identical levels of expression in both taxa. It is possible that low levels of nocturnal CO_2_ fixation via PEPC evolved in the ancestor of both species for non-photosynthetic reasons; for example, tobacco leaves (Kunitake et al., 1959) and cotton ovules (Dhindsa et al., 1975) show various levels of carbon concentration, and fixation of cytosolic CO_2_ via PEPC is required to replenish the citric acid cycle (Aubry et al., 2011). Because these pathways already exist, slight upregulation of some components in a shared ancestor may have enabled the origins of CAM in certain lineages (Bräutigam et al., 2017).

However, gene expression alone clearly does not make a species CAM, and further refinement is necessary. Refinement of CAM may take the form of improving secondary metabolic processes or genomic characteristics that allow for strong and constitutive CAM to exist. For example, fine tuning of carbohydrate turnover is necessary for CAM function (Borland et al., 2016; Ceusters et al., 2014). Experiments placing *Mesembryanthemum crystallinum* in CO_2_-free air at night resulted in a dampened CAM cycle (Dodd et al., 2003) and comparative RNA-seq in facultative CAM and C_3_/CAM comparisons has shown increased reliance on carbohydrate breakdown as CAM function increases (Brilhaus et al., 2016; Heyduk et al., 2018a). Circadian regulation of CAM genes in pineapple was shown to be the result of promoters that induce evening expression (Ming et al., 2015), and genes that link the clock to metabolic outputs had shifts in expression phasing between *Agave* (CAM) and *Arabidopsis* (C_3_) (Yin et al., 2018). Variation in timing and magnitude of expression of various light sensing and clock genes has been described not only here in *Erycina*, but also in *Agave* (Abraham et al., 2016). In all, the growing evidence suggests that the secondary processes that make a CAM plant, including stomatal regulation, light sensing and downstream transcriptional responses, and carbohydrate metabolism feedbacks, undergo fine-tuning along the evolutionary trajectory between C_3_ and CAM. Further work that characterizes closely related C_3_ and CAM species has the potential to greatly advance our understanding of the integration of nighttime CO_2_ acquisition with more complex regulatory pathways.

## Data Availability

Raw sequence reads are available on NCBI Short Read Archive, under the BioProject PRJNA483943. All scripts used that are not part of existing programs are available at www.github.com/kheyduk.

## Acknowledgements

The authors would like to sincerely thank Orquideas Tropicales in Panama for generously donating *Erycina crista-galli* plants. Additional gratitude goes to Saravanaraj Ayyampalayam for assistance with data processing, Jeremy Ray for assistance with RNA preparation, and Rishi Masalia for giving feedback on figures. This work was supported by NSF grants to J.L.M, V.A, and K.S (DEB #1442199 and 1442190) and the by Smithsonian Tropical Research Institute.

## Author Contributions Statement

JLM and VAA conceived and led the project; KS and KW collected, phenotyped plants, and conducted continuous gas exchange measurements, and KS additionally sampled *E. crista-galli* for RNA-seq; VAA and TL grew and sampled *Erycina pusilla* individuals for RNA and physiology; KH and MH sequenced and analyzed the data and prepared the manuscript; JLM oversaw general experimental design, and all authors contributed to the final version of this manuscript.

